# Gene Drives Across Engineered Fitness Valleys: Modeling a Design to Prevent Drive Spillover

**DOI:** 10.1101/2020.10.29.360404

**Authors:** Frederik J.H. de Haas, Sarah P. Otto

## Abstract

Engineered gene drive techniques for population replacement and/or suppression have potential for tackling complex challenges, including reducing the spread of diseases and invasive species. Unfortunately, the self-propelled behavior of drives can lead to the spread of transgenic elements beyond the target population, which is concerning. Gene drive systems with a low threshold frequency for invasion, such as homing-based gene drive systems, require initially few transgenic individuals to spread and are therefore easy to implement. However their ease of spread presents a double-edged sword; their low threshold makes these drives much more susceptible to spread outside of the target population (spillover). We model a proposed drive system that transitions in time from a low threshold drive system (homing-based gene drive) to a high threshold drive system (underdominance) using daisy chain technology. This combination leads to a spatially restricted drive strategy, while maintaining an attainable release threshold. We develop and analyze a discrete-time model as proof of concept and find that this technique effectively generates stable local population suppression, while preventing the spread of transgenic elements beyond the target population under biologically realistic parameters.

## 2 Notes from reviewers

## 3 Introduction

Engineered gene drives are self-replicating genetic constructs that can bias transmission of desired alleles (usually called payload genes) to progeny, allowing them to rapidly increase in frequency even when the alleles are selected against [Burt, 2003]. Introgression of these transgenic genes into sexually reproducing species could potentially solve many pressing environmental and humanitarian problems, ranging from public health and agriculture to conservation [National Academies of Sciences et al., 2016]. Examples include modifications of the mosquito genome to reduce its capacity to serve as a disease vector (e.g. reducing the number of female mosquitoes, their longevity or their ability to support development and transmission of the pathogen) [Isaacs et al., 2012]. Other examples include driving genes that protect species at risk by eradicating invasive species or reducing pest damage in agriculture by replacing resistant alleles with their ancestral equivalents to restore vulnerability to pesticide or herbicide [National Academies of Sciences et al., 2016].

There are two general approaches to gene drive: population suppression and population alteration. Population suppression makes use of gene drive systems to purposefully sterilise or kill the target population while population alteration uses gene drive systems to alter the genomic composition without the aim to sterilise or kill the target population. A successful example of population suppression in the laboratory was shown by Kyrou et al. [2018] for *Anopheles gambiae*. They used a CRISPR-Cas9 gene drive targeting a doublesex gene (Agdsx) that lead to a bias in the sex ratio by rendering homozygous females completely sterile. Within 7–11 generations egg production was reduced to the point of total population collapse.

Extreme caution with population suppression is required as it could send a ripple through ecosystems, endangering many other plants and animals than just the target species. For example, mosquitoes that live in the arctic of Canada & Russia fly around in thick swarms and make up a huge part of the biomass there. They pollinate arctic plants and are a major food source for migrating birds so their loss could be disastrous to the ecosystem [Thien, 1969]. A much less invasive approach to gene drive is population alteration. This approach potentially has less ecological harm, but it might require multiple drives for each disease carried by the vector. Isaacs et al. [2012] discovered that *Anopheles stephensi* mosquitoes expressing m1C3, m4B7, or m2A10 single-chain antibodies (scFvs) have significantly lower levels of infection compared to controls when challenged with *Plasmodium falciparum*, a human malaria pathogen. Further research needs to show if expression of a single copy of a dual scFv transgene can completely inhibit parasite development, potentially without a fitness cost on the mosquito.

Although gene drives hold much promise, they have the strong possibility of unwanted ecological effects. Major risks include the unintended spread of the payload beyond the area of interest due to unwanted migration of individuals and the spread of the payload beyond the target species due to hybridization. Low threshold gene drive systems such as homing-based endonuclease gene drive (HEG) are especially susceptible to such spillovers. These systems exploit homology directed repair (HDR) to replace a targeted, naturally occurring genetic sequence with an engineered construct. Homing-based gene drives work by transcribing an endonuclease (often Cas9) and a guide RNA (gRNA) in the germline. The endonuclease then triggers a double stranded break at the complementary site of the gRNA which can be repaired by either one of two pathways: non-homologous end joining (NHEJ) or homologous end joining (HEJ). In NHEJ the break ends are directly ligated without the use of a homologous template. This often leads to mutations arising in the process, preventing the gRNA from recognizing the sequence for future double stranded breaks. However, when the double stranded break is repaired by HEJ, which requires the homologous chromosome carrying the construct, the result is a copy of the homologous chromosome at the double stranded break: creating effectively homozygous cells from heterozygous cells. The result is equivalent to meiotic drive, *δ*, leading to a greater fraction of gametes carrying the construct than expected under Mendelian inheritance (specifically, heterozygotes produce a fraction 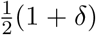 of gametes carrying the driven allele).

High conversion rates have already been achieved in the lab by the CRISPR-Cas9 system for yeast (*δ* > 0.995, [DiCarlo et al., 2015]), fruit flies (*δ* > 0.97, [Gantz and Bier, 2015]), and the malaria vector mosquito, *Anopheles stephensi*, with engineered malaria resistance (*δ* > 0.98, [Gantz et al., 2015]). These drives require few transgenic individuals for gene drive to occur (low threshold), making them very capable of spreading between subpopulations following introduction by migration or following hybridization between species [Marshall, 2009]. Numerous countermeasures have been proposed to reverse gene drives such as synthetic CRISPR-cas9 based reversal drives like CATCHA [Wu et al., 2016] and ERACR [Gantz and Bier, 2016]. However, reversal drives are techniques to fully eliminate rather then locally confine the drive, which is often the desired outcome.

To address the issue of local confinement, Noble et al. [2019] analyzed a new self-exhausting drive technique that relies on CRISPR-mediated homing-based gene drive called a “daisy-chain drive”. The daisy-chain drive separates multiple interdependent homing-based drive components across the organism’s genome, each bearing a cost (drive load). Each genetic element drives the next in the chain. The final element carries the payload and is driven to higher and higher frequencies due to the preceding drive elements. Because no element can drive itself and the first element is not driven at all, the eventual outcome is that all elements will be lost from the population over time due to natural selection against the drive load, and the population returns to its original wildtype state. Unfortunately, as the drive disappears, so does the payload gene, rendering this system ineffective for a persistent payload introgression.

Persistent and localized introgression would be possible if the daisy chain alleles drive a high threshold drive system (such as underdominance) past its high threshold frequency. The advantage of such a construct is that it would be eliminated from adjacent populations once the daisy chain is exhausted due to their low frequency (below the threshold). This design was first proposed conceptually by Min et al., under the name “daisy quorum drive” ([Min et al., 2017]) but has not yet been modeled, which is our goal here. This new drive design is predicted to spread through a population until all of its daisy elements have been lost, at which point its fitness becomes frequency-dependent. The result is an engineered population surrounded by wildtype populations with limited mixing at the boundaries due to the reduced fitness of crosses between wildtype individuals and individuals carrying the engineered alleles.

Drives that exhibit a high invasion threshold, such as engineered underdominance (EU) and Wolbachia drives, provide a natural means to achieve localization. These drives exhibit frequency-dependent dynamics where the drive can only spread if its frequency exceeds the invasion threshold. Below this level, the frequency of the drive will decrease, leading to its loss from the population. Spread of a threshold drive across a patchy environment is more difficult and becomes highly unlikely or even impossible when the invasion threshold is 50 percent or higher [Barton and Turelli, 2011]. Threshold gene drive mechanisms were among the first forms of gene drive proposed and the only one to be deployed in the wild [Buchman et al., 2018]. Because organisms that migrate to other populations will be in the minority and consequently are selected against, underdominance with high thresholds poses a substantially lower risk of unintended spread.

An ideal gene drive system to alter wild populations would exclusively affect organisms within the boundaries of consenting communities. Here we present a simple population genetics model to elucidate the evolutionary dynamics of a new design that meets this criterion by combining daisy-chain drive with a two-locus underdominance system (Figure 1). The daisy-chain drive system provides a short window of time to drive the frequency-dependent underdominance system before it disappears, effectively transitioning from a low to a high threshold control system. By exploring spatial models in the face of recurrent migration, we find theoretical support that this new design could reduce the risk of gene drive spillovers while providing persistent payload introgression to the target population(s).

**Figure 1:**
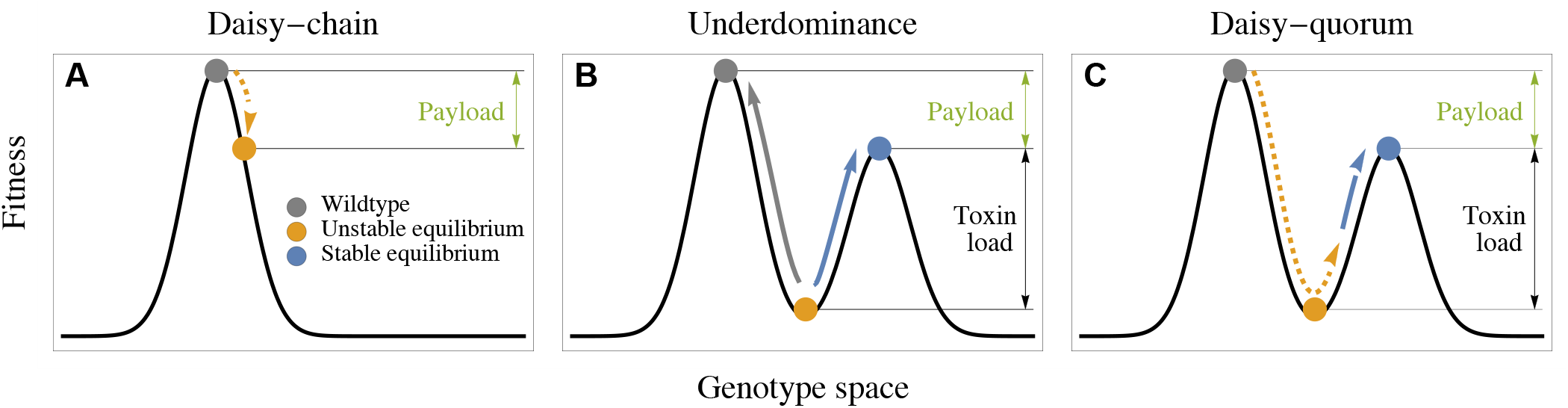
Conceptual overview of the different control systems. (A) A daisy-chain drive introgresses a payload temporarily, ultimately reverting back to the wildtype state (grey). (B) The dynamics of an underdominance gene drive is frequency dependent: left of the valley the population is driven back to a wildtype composition, while right of the valley it evolves toward the lower peak expressing the payload. (C) The proposed design (daisy-quorum drive) includes both systems and uses the temporary low-threshold daisy chain to drive an underdominant construct past the valley, after which it evolves toward the other peak with the desired payload.

## 4 Methods and results

We formulate a deterministic model that considers the evolution of a large population of diploid organisms. We first introduce the model for a single population and then build up to an array of populations under recurrent migration to explore the spatial spread of the engineered construct. We refer to the loci in our model by bold letters (e.g. **A**) and to the alleles by the italic letters (e.g. *A* or *a*)

### 4.1 Drive design

We model a multilocus bi-allelic population with random mating using discrete-time dynamics. Our proposed drive system consists of two components. The first component encodes a daisy-chain gene drive, which can be of arbitrary length, and the second component is a two-locus engineered underdominance system with an added payload gene on each transgenic allele. We explore here a four-locus genetic architecture as an example case (Table 1), although the daisy chain could contain more loci if a longer acting drive is needed [Min et al., 2017] [Noble et al., 2019]. Unless otherwise stated, we assume that the loci involved are freely recombining 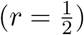 and that all dynamical systems are initialized at a proportion 1 − *f*_0_ of *abcd* gametes and *f*_0_ of *ABCD* gametes.

**Table 1:** Four-locus daisy-quorum drive. Loci **A** and **B** are part of the daisy-chain component and loci **C** and **D** make up the two-locus underdominance component. The daisy-chain transgenic alleles (**A** and **B**) have a fitness cost of *s*_*d*_ (drive load) and the engineered underdominance transgenic alleles have a toxin-load *s*_*t*_ and a payload *s*_*p*_, which interact according to Table 2. The gRNA of locus **A** is complementary to the wildtype allele *b*, and locus **B** carries two gRNAs to cut the wildtype alleles *c* and *d* at the underdominance loci.

| Daisy-chain Component |  |  |  | Underdominance Component |  |  |  |
| --- | --- | --- | --- | --- | --- | --- | --- |
| Locus A |  | Locus B |  | Locus C |  | Locus D |  |
| *a* | Wildtype | *b* | Wildtype | *c* | Wildtype | *d* | Wildtype |
| *A* | CRISPR-Cas9<br>gRNA targetting *b*<br>**Drive load** ($s_d$) | *B* | CRISPR-Cas9<br>gRNA targetting *c*<br>gRNA targetting *d*<br>**Drive load** ($s_d$) | *C* | **Payload** ($s_p$)<br>Suppressor of *D*<br>Promoter<br>**Toxin load** ($s_t$) | *D* | **Payload** ($s_p$)<br>Suppressor of *C*<br>Promoter<br>**Toxin load** ($s_t$) |

The first component (daisy-chain) consists of a linear series of *n* loci arranged such that each element drives the next in the chain. The final element in the chain drives the “cargo”, which in our model are the two underdominance loci **C** and **D**. Each transgenic allele of the daisy chain consists of a CRISPR-cas9 complex and a drive load, which bears a fitness cost of *s*_*d*_. The drive load helps ensure that all CRISPR-cas9 elements are counterselected and eventually disappear from the system.

In the case of the daisy-chain, the gRNA at locus *i* in the chain guides the Cas9 nuclease to the wildtype at locus *i* + 1, inducing a double stranded break, which will be repaired by homologous recombination with probability *δ*. We do not incorporate natural resistance alleles to the daisy-drive elements because their effects are minimal due to the transient nature of the daisy chain drive (see supplementary material).

The two-locus underdominance component is structured following the design proposed by Davis et al., [Davis et al., 2001]. The two transgenic alleles consist of four elements each: a payload, a suppressor, a promoter and a toxin. The two engineered alleles are carried on separate non-homologous pairs of chromosomes, each of which produces a toxin unless the suppressor on the other transgenic allele is also present. Specifically, in the absence of the matching suppressor, the promoter will cause the toxin gene to be expressed, while individuals who carry one or more copies of both constructs are viable because both suppressors are present in the genome and neither toxin gene is expressed (Table 2). This scheme creates a frequency-dependent fitness regime, promoting the state of carrying both engineered constructs or none and selecting against individuals carrying one but not the other transgenic allele.

**Table 2:** Relative fitness for each diploid genotype in the two-locus underdominant component. The toxin *s*_*t*_ is expressed in individuals carrying either a *C* or *D* but not both. The payload *s*_*p*_ is expressed if the individual carries at least 1 copy (dominantly expressed). Both the toxin and the payload affect individual fitness multiplicatively.

| | $cd$ | $cD$ | $Cd$ | $CD$ |
| --- | --- | --- | --- | --- |
| $cd$ | 1 | $(1 - s_t)(1 - s_p)$ | $(1 - s_t)(1 - s_p)$ | $(1 - s_p)$ |
| $cD$ | $(1 - s_t)(1 - s_p)$ | $(1 - s_t)(1 - s_p)$ | $(1 - s_p)$ | $(1 - s_p)$ |
| $Cd$ | $(1 - s_t)(1 - s_p)$ | $(1 - s_p)$ | $(1 - s_t)(1 - s_p)$ | $(1 - s_p)$ |
| $CD$ | $(1 - s_p)$ | $(1 - s_p)$ | $(1 - s_p)$ | $(1 - s_p)$ |

The effects on fitness (toxin load *s*_*t*_, drive load *s*_*d*_ and payload *s*_*p*_) are assumed to determine individual fitness multiplicatively. Table 2 illustrates the effect of locus **C** and **D** on individual fitness where the payload (*s*_*p*_) is dominantly expressed. In the supplementary material we explore the effect of different payload fitness regimes: multiplicative dominance and a recessive case. To derive the relative fitness for the full genotype, the fitness of the **CD** genotype is multiply by (1 − *s*_*d*_)^*ρ*^ where *ρ* equals the number of *A* and *B* alleles.

### 4.2 Dynamics in an isolated population

The recursion equations for the 16 (2^4^) gamete types are too elaborate so we report them in the supplementary *Mathematica* file. Instead we focus on the dynamical equations of the first and second component in isolation (Table 1). We use recursion equations to take the population through diploid selection followed by meiosis with drive and recombination.

Assuming random mating in a single isolated population, the recursion equations for the frequency *X*_*ij*_ of the gamete *ij* at loci **A** and **B** are:

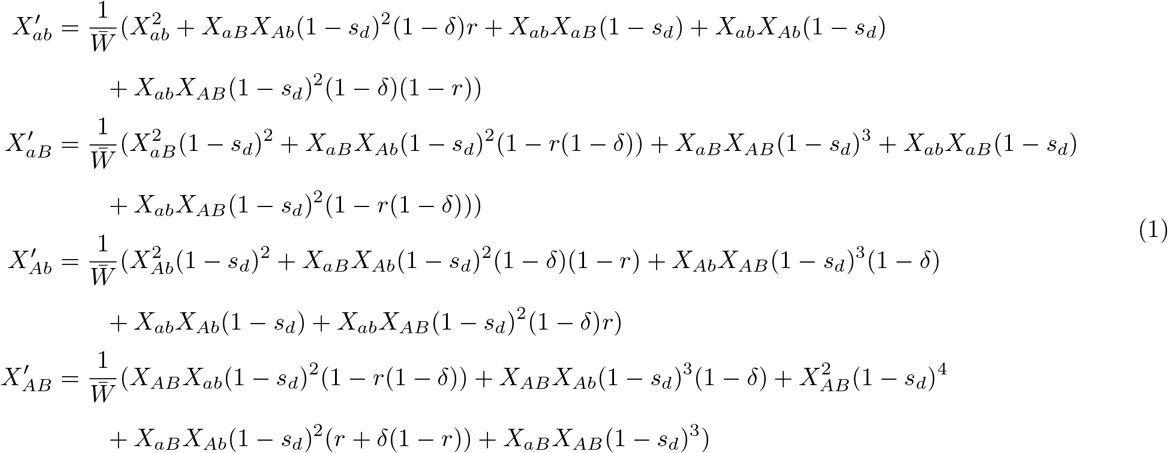

where *X*_*ab*_ + *X*_*aB*_ + *X*_*Ab*_ + *X*_*AB*_ = 1 and 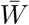 is the average fitness. In the supplementary material we add a mating table for these recursion equations (Table S1). An automated algorithm for deriving the dynamical equations for longer daisy chains is provided in the supplementary *Mathematica* file.

The recursion equations for the second underdominance component, describing the frequency *X*_*ij*_ of the gamete *ij* at loci **C** and **D**, are:

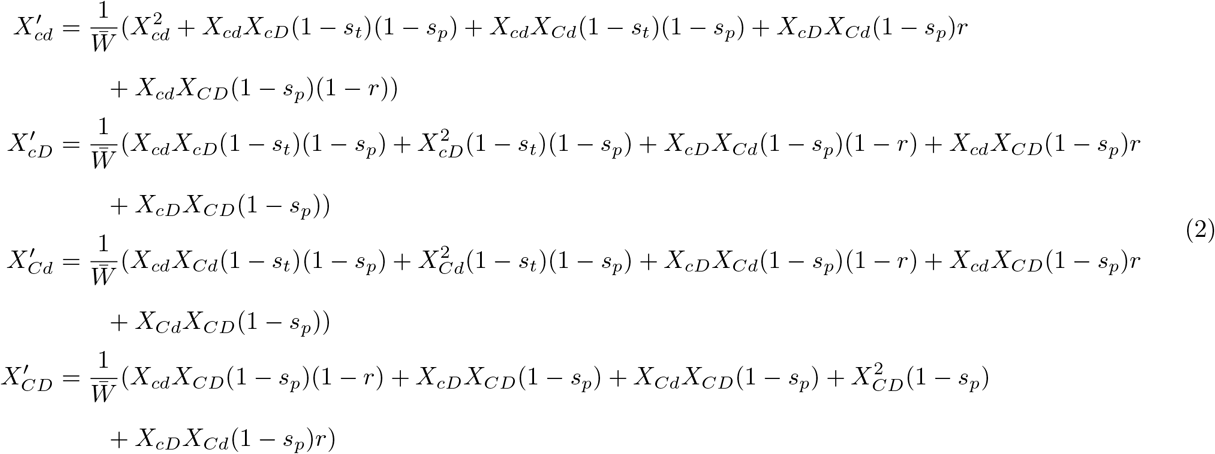

where *X*_*cd*_ + *X*_*cD*_ + *X*_*Cd*_ + *X*_*CD*_ = 1 and 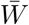 is the average fitness (the sum of the numerators).

We assume that *X*_*cD*_ = *X*_*Cd*_ at *t* = 0, which implies that their frequencies will remain the same at all time points in the future (Equation 2). By inspection of Equation 2, *X*_*cd*_ = 1 and *X*_*CD*_ = 1 are fixed point solutions to the underdominant component of the dynamical equations and a local stability analysis indicates that both are locally stable. A third fixed point represents the co-existence of alleles *C* and *c*, as well as *D* and *d*, and is locally unstable (see Supplementary Material for ternary plots of alternative payload fitness regimes). In Figure 2, we illustrate the location of the stable equilibria (blue and grey vertices) and the separatrix (the boundary separating two basins of attraction in a dynamical model) for different payloads *s*_*p*_ when the toxin is relatively ineffective (*s*_*t*_ = 0.1) or fully effective (*s*_*t*_ = 1.0). The separatrix is calculated numerically. Example dynamics confirm that the system converges to either fixed point depending on whether initialized to the left or right side of the separatrix.

**Figure 2:**
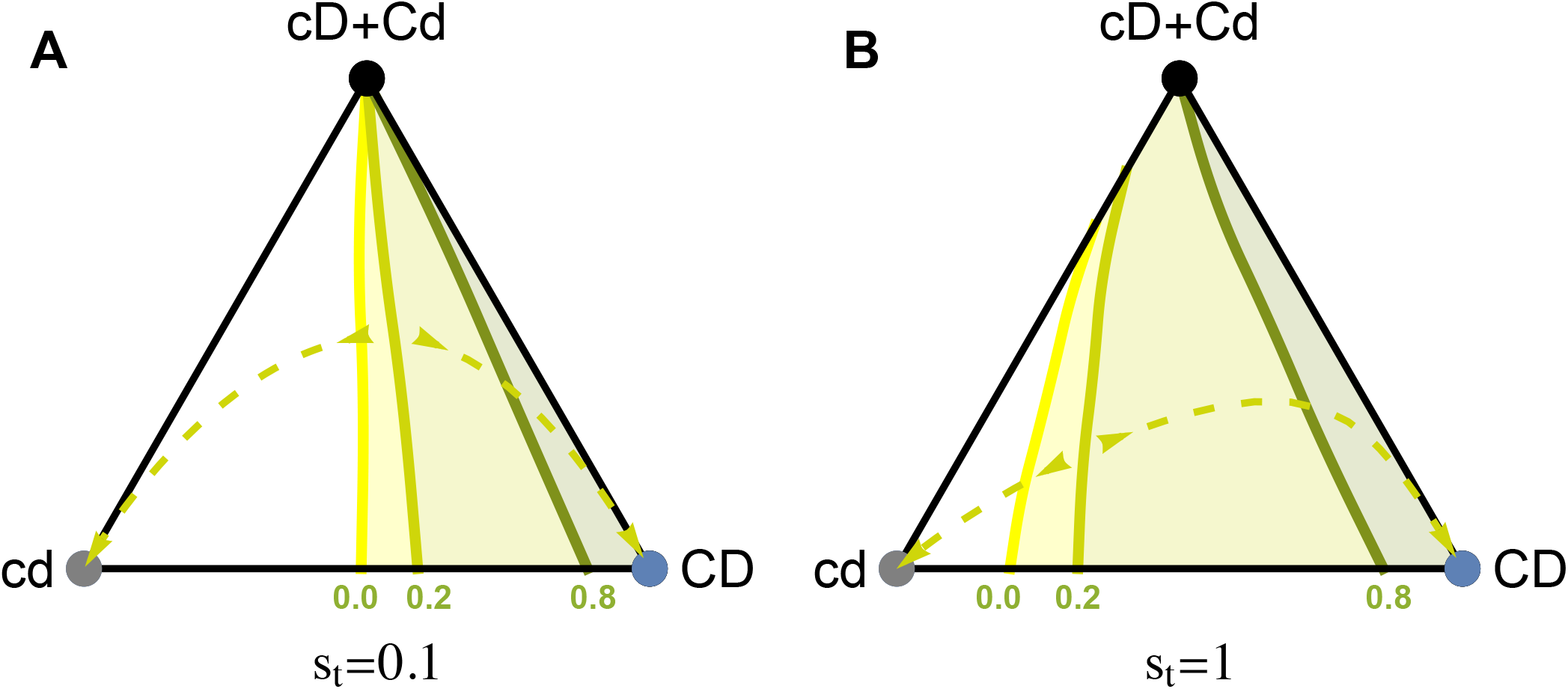
Ternary plot illustrating the separatrix (the boundary separating basins of attraction to different fixed points) for the two-locus underdominance construct with a toxin load of (A) *s*_*t*_ = 0.1 or (B) *s*_*t*_ = 1.0 and a payload of *s*_*p*_ = 0.0, 0.2 and 0.8 (green curves). Locus **C** and **D** are recombining freely 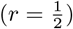. The dashed curves are example dynamics for *s*_*p*_ = 0.2.

As the payload increases (larger *s*_*p*_), the separatrix moves to the right in Figure 2, reducing the basin of attraction of the engineered underdominance construct (haplotype *CD*). Higher payloads thus make it harder to drive such constructs into a population, but they also provide stronger fitness effects if the construct is able to fix. By contrast, as the toxin load increases (larger *s*_*t*_), the basin of attraction to *CD* increases. While somewhat counterintuitive, this occurs because the toxin load can be suppressed by even one copy of the alternate allele, making haplotypes *Cd* and *cD* strongly selected against when *cd* is common (the wildtype) but only weakly selected against when CD is common, causing the fitness surface to fall faster near *cd* but not near *CD* when the load is stronger.

To prevent unintentional spread into neighbouring populations, it is desirable to design a drive where the separatrix is shifted far towards the transgenic haplotype so invasion will only occur once a significant number of transgenic haplotypes have been introduced into a new population. However, shifting the unstable equilibrium too far to the transgenic haplotype will cause the population to revert to the wildtype state more easily due to drift and incoming wildtype migrants from neighbouring populations. We therefore suggest that a robust system of localization is designed such that the unstable equilibrium is near the center of Figure 2, which can be achieved by adjusting the toxin load *s*_*t*_ relative to the payload *s*_*p*_.

To derive the dynamics for all gamete types, the two components are linked by allele *B* driving allele *C* and *D*, just like allele *A* drives *B*.

### 4.3 Invasion analysis

We next perform an invasion analysis to determine the drive rate *δ*_*c*_ needed for the transgenic alleles *C* and *D* to spread. Here *δ*_*c*_ is treated as a constant value, equalling the frequency of individuals carrying the last daisy-chain locus 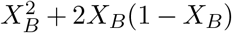 times the drive strength *δ*. The conditions for the spread of the cargo when rare are that the following condition holds (assuming 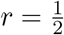 and *B*, *C* and *D* are unlinked):

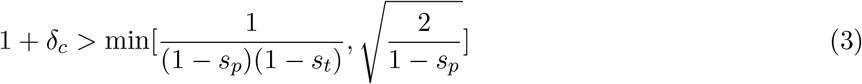

The minimum term describes the conditions under which either allele *C* or *D* could spread on their own (first criterion) or the combined *CD* haplotype could spread (second criterion). With a low toxin load (*s*_*t*_), the first criterion is easier to satisfy, and we expect the alleles to spread individually. With a high toxin load, however, only the combined haplotype *CD* will spread, as long as drive is strong enough to counter the haplotype being broken down by recombination.

Given that 0 ≤ *δ*_*c*_ ≤ 1, a two-locus underdominant construct carrying a dominant payload with *s*_*p*_ > 0.5 can never invade, no matter the strength of the drive. With multiplicative dominance or recessive payload fitness regimes, a higher payload can be invaded (see Supplementary Material), however an interior basin of attraction prevents the payload from reaching fixation under these fitness regimes.

Figure 3 illustrates the contrasting effects of a constant “cargo” load compared to a two-locus underdominant “cargo” for different initial release frequencies *f*_0_. A constant cargo load is modeled as a third locus *C’* that only contains a payload, as is generally considered with daisy chain drives. Such a constraint cargo does not spread the payload to fixation as it is always being selected against while the underdominant system allows the fixation of the payload provided that it is released at a high enough initial release frequency. Due to the daisy-chain drive, an underdominance construct carrying a payload of *s*_*p*_ = 0.2 can spread at a far lower initial release frequency than predicted for the underdominance component on its own (threshold of 0.24 when introducing *CD*, Figure 2). Panel D illustrates the scenario for a lower initial release frequency where the daisy drive proved to be not powerful enough to push the underdominance construct past the separatrix of Figure 2.

**Figure 3:**
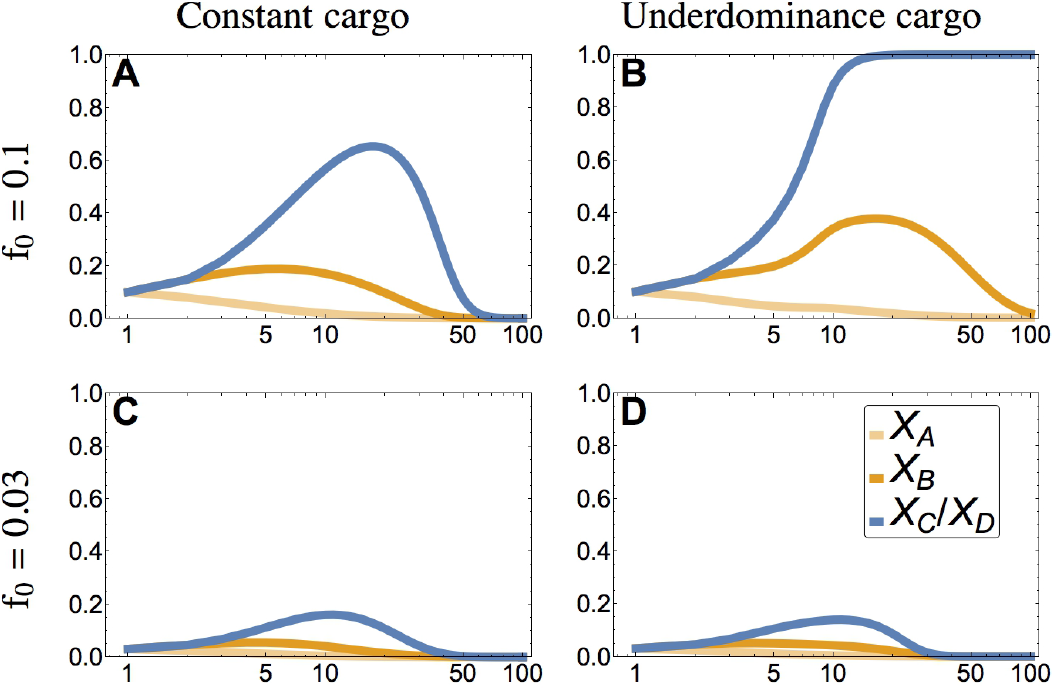
Top row illustrates dynamics of a daisy chain pushing a constant payload *s*_*p*_ = 0.15 while bottom row shows the daisy chain pushing an underdominance construct, also with a payload of *s*_*p*_ = 0.15. The panels are initialized with gametes *ABCD* at frequency *f*_0_ = 0.1 (left) and *f*_0_ = 0.03 (right). Other parameters are: *s*_*t*_ = 0.9, *d* = 0.95, *s*_*d*_ = 0.05 and 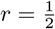.

### 4.4 Spatial spread of the gene drive

We next model a stepping-stone model of *M* interconnected populations with migration rate *m* between adjacent patches. Migration is followed by dynamics within each path, as described in section 2.2. The frequency of gametes of type *i* after migration for a non-boundary population *j* is given by: 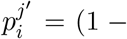 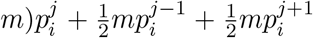. The boundary populations only give and receive migrants from the interior: 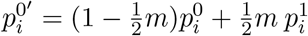 and 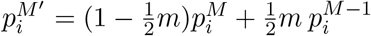.

Migration rate *m* is a crucial parameter to determine the spread of the payload through interconnected populations. In Figure 4 we show that our gene drive construct can be locally maintained in the face of recurrent migration and yet remain spatially restricted because migration into neighbouring demes does not introduce enough of the daisy chain to reach the threshold for invasion. Figure 4 (panel B) shows the range of migration rates where the construct will remain spatially restricted (*m* < 10^−1.3^). For increasing migration rates, the payload spreads further (e.g., to patches +1 and −1 with *m* = 0.05) until migration rate reaches a critical value above which the drive construct is swamped within the initial population and fails to spread. The payload frequency shows an almost stepwise function at the boundary, from a frequency of 1 to a frequency of 0 (gray zone).

**Figure 4:**
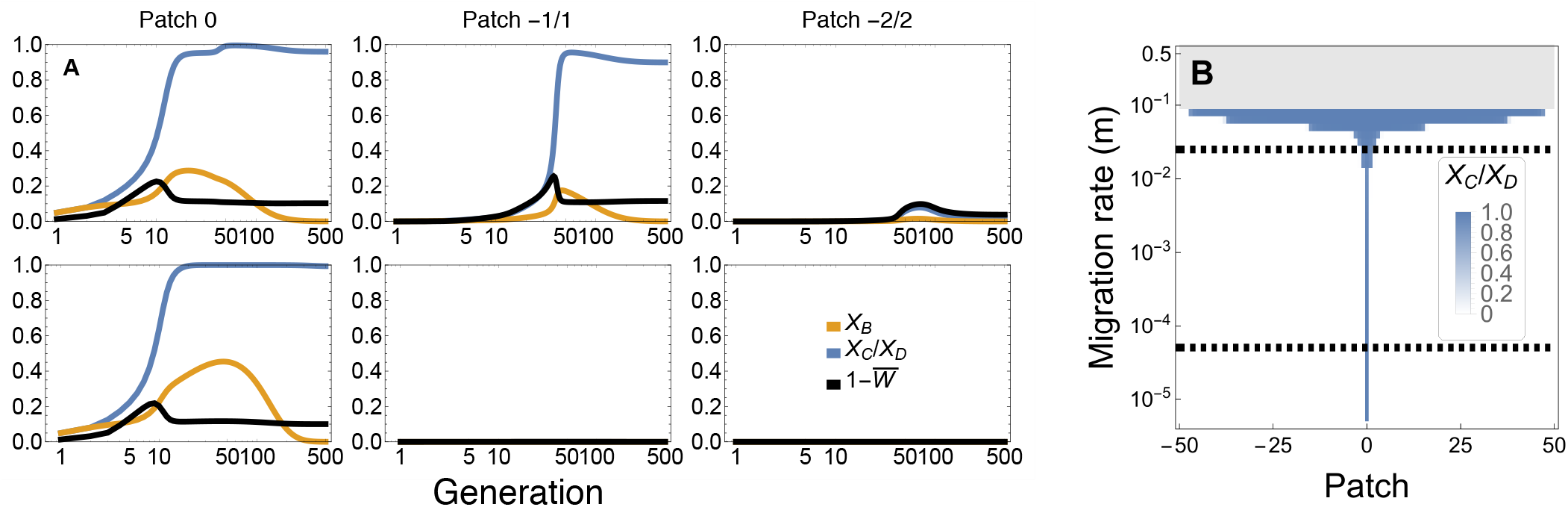
Dynamics of spread of a payload for different migration rates when *ABCD* gametes are introduced into the center patch (patch 0) at an initial frequency of *f*_0_ = 0.05. Top and bottom row of panel (A) show 3 interconnected patches and their dynamics for a migration rate of *m* = 0.05 and *m* = 0.00005, respectively (see Table 3 for other parameters). For a relatively high migration rate (top row), spread to neighbouring populations is more likely than for lower migration rates (bottom row). As predicted, the drive disappears from the system due to the drive load *s*_*d*_ = 0.02 (orange curves). Panel (B) shows the frequency of the payload alleles, *X*_*C*_ or *X*_*D*_, at 1000 generations (blue shading) across all patches (x axis) for a variety of migration rates (y axis). In the gray area of panel (B), migration into the center patch (patch 0) is high enough to swamp the engineered construct causing no spread to occur anywhere.

Given a particular migration rate *m* within a given natural system, the power of the daisychain combined with the location of the separatrix determine how far the drive can spread through neighbouring populations. Parameters that affect the daisychain are the drive load *s*_*d*_, the drive rate *δ*, and the length of the daisy chain *n*. Parameters affecting the separatrix are the payload *s*_*p*_ and the toxin load *s*_*t*_. These parameters could be adjusted during the engineering of the construct, affecting the risk of spread as shown in Figure 5.

**Table 3:**
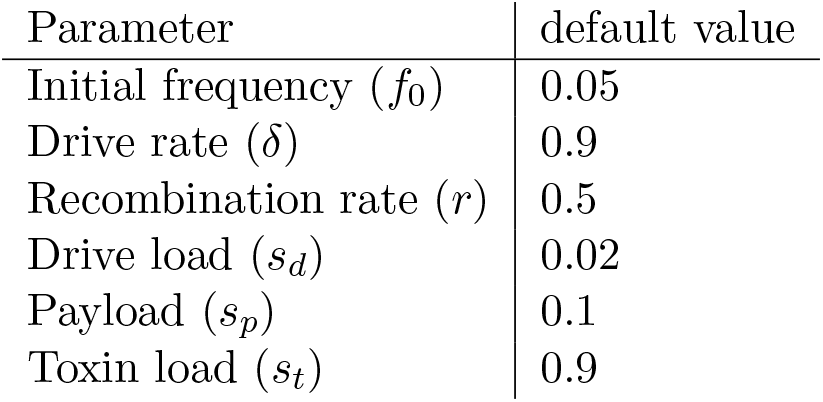
Default parameter values for Figure 4 and 5

**Figure 5:**
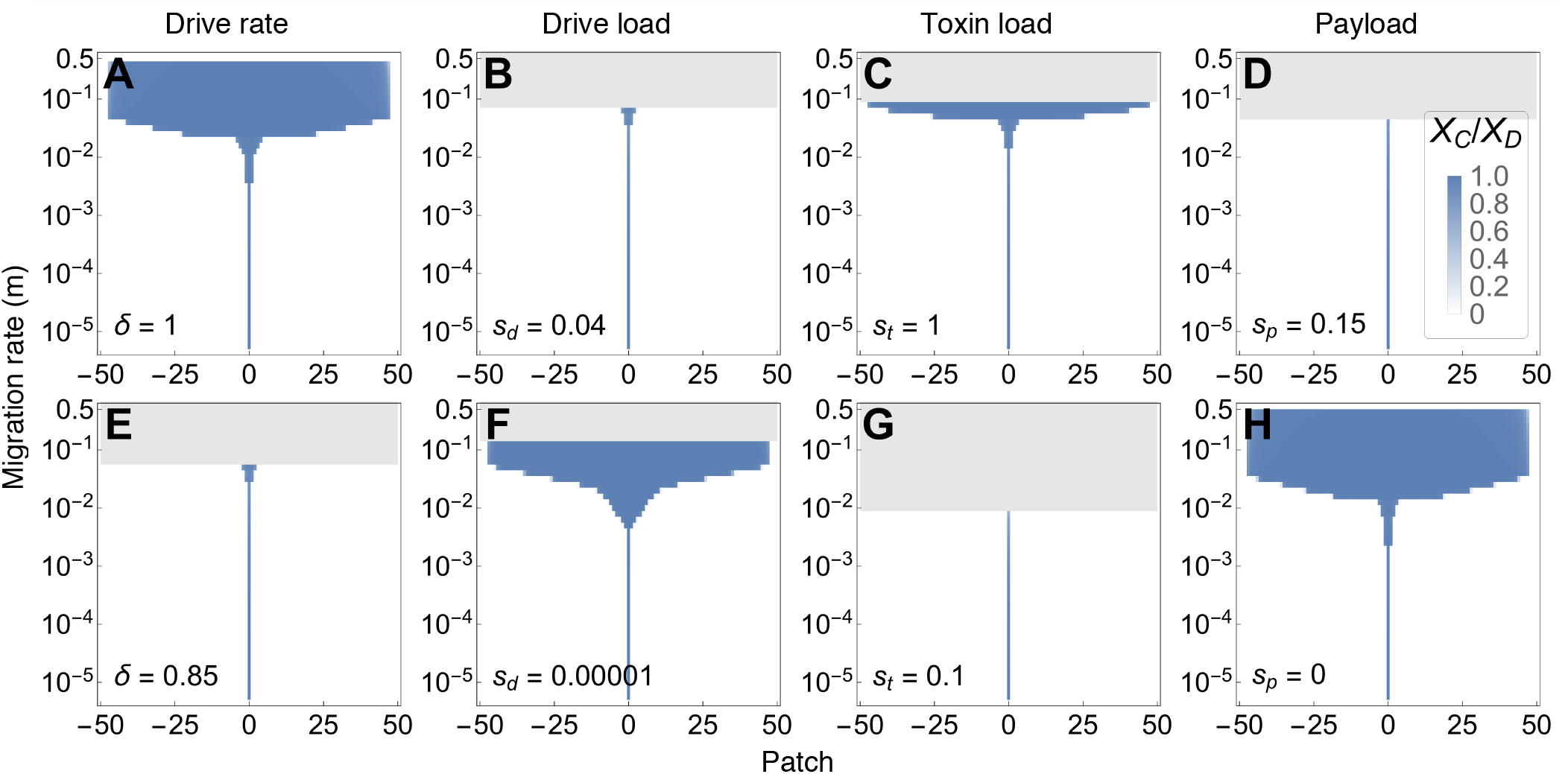
Illustration of the effect of chancing a single parameter on the results shown in Figure 4 (panel B) for the default parameters (Table 3). The frequency of the payload is shown in blue after 1000 generations when *ABCD* gametes are introduced into the center patch (patch 0) at frequency *f*_0_ = 0.05. Patches are linearly connected and exchange migrants every generation at a rate *m*. Subfigures from left to right change the following single parameter up (top figures) or down (bottom figures): drive rate *δ*, drive load *s*_*d*_, toxin load *s*_*t*_ and the payload *s*_*p*_

Lowering the drive load *s*_*d*_, increasing the drive rate *δ* or lengthening the daisy chain *n* (see supplementary material) results in a daisy chain that persists for longer and spreads the underdominant construct to more populations. Interestingly, a decrease in the toxin-load reduces the risk of spread of the payload to neighbouring populations, even under high migration rates. This is because the lower the toxin load, the closer the separatrix becomes to the state where the underdominant construct *CD* is fixed (Figure 3). Thus, somewhat counterintuitively, a lower toxin load provides a higher degree of safety in isolating the underdominance construct from spreading into neighbouring patches once the daisy-chain drive has exhausted itself.

Stochastic simulations confirm that these results are robust to finite population sizes in cases where the population persists (Figure S4). In cases where the payload drives the local population extinct, recolonization eventually occurs from neighbouring populations (Figure S5).

## 5 Discussion

Low threshold gene drives such as homing-based gene drives risk spillover to neighbouring populations due to migrating individuals. The gene drive design that we study here transitions in time from a low threshold drive (homing-based gene drive), which is especially susceptible to these spillovers, to a high threshold drive (two-locus underdominance), which is more resilient to spillover. This transition is achieved by using a daisy-chain technique recently developed by Noble et al. [2019]. We successfully show that this design prevents spread of the payload under biologically realistic scenarios (Figures 4 + 5).

Previous models of homing-based gene drives have focused on the rise of resistance alleles that would prevent the drive from completely reaching fixation in the target population and ultimately revert the system back to the wildtype state. Here we do not explicitly take resistance alleles into account because the daisy-chain drive system, by its nature, decreases in effectiveness. Adding resistance alleles does not drastically change the dynamics of our results (see Supplementary Material). However, we note that the two-locus underdominance construct is susceptible to mutations that decrease or compensate for the payload, which would affect the long-term persistence of the engineered suppression. By contrast mutations that reduce the toxin-load have little effect because the toxin-load is transient and disappears once the underdominance constructs (loci *C* and *D*) are fixed. Unlike Dhole et al. [2018], we recommend that both transgenic alleles *C* and *D* carry the payload *s*_*p*_ which might better resist the evolution of a compensatory mutation within the payload genotype due to the existence of multiple gene copies.

We also tried an advantage to population alteration, compared to population suppression, that emerges in localized drive systems like the one explored here. If the payload causes local populations to decline to extinction (see Figure S5), wildtype individuals from neighbouring patches will eventually recolonize and return the system to its original state. By contrast, strategies that involve population alteration persist for longer because migrants continue to face competition and toxic interactions with individuals in the target population carrying the underdominance construct (see Figure S4). Thus the engineered construct investigated here are most suited for population alteration, rather than population suppression, when a persistent and localized effect is sought.

The possibility that gene drive systems provide to alter the genome of species is concerning to many people. Implementing gene drives, accidentally or on purpose, can have long lasting effects on biological systems. It is clear that gene drives have the potential to solve many ecological challenges, however, all consequences of triggering a gene drive cannot be foreseen. Models like the one we present here are important first steps, but they can only be used as a rough guide, while much more experiments and refined models are required before release of gene drives in the wild [National Academies of Sciences et al., 2016].

In summary, we modeled a potential solution to the spillover and persistence problems posed by gene drive systems under realistic biological parameters using a combination of homing based gene drive and two-locus underdominance. Lab-based research is needed to determine the feasibility of developing an underdominant daisy-quorum drive. In particular, controlled experiments are needed to determine the likelihood of mutations reversing the payload (reducing persistence) and the risk of unintended side effects, such as self-driving elements arising from the daisy-chain genes (increasing spillover risk).

## 6 Acknowledgements

We would like to thank the Otto lab for helpful feedback and for their valuable input to this project. This work was supported by a Discovery grant from the Natural Sciences and Engineering Research Council of Canada to SPO (NSERC RGPIN-2016-03711) and stipend support to FJHdH from an NSERC CREATE grant (BIOS2).

## 7 Supplementary Material

### 7.1 Daisy-chain length

The length of the daisy chain is an important determinant of the maximum attainable frequency of the payload. Here we illustrate this effect for daisy chains of different lengths by tracking the maximal frequency attained by the last element in the chain. When the number of loci involved in the daisy chain increases, there is a stronger and longer force pushing the last element to a higher frequency (Figure S1). The inline figure shows an example of the dynamics of a daisy chain system for 3 loci.

**Figure S1:**
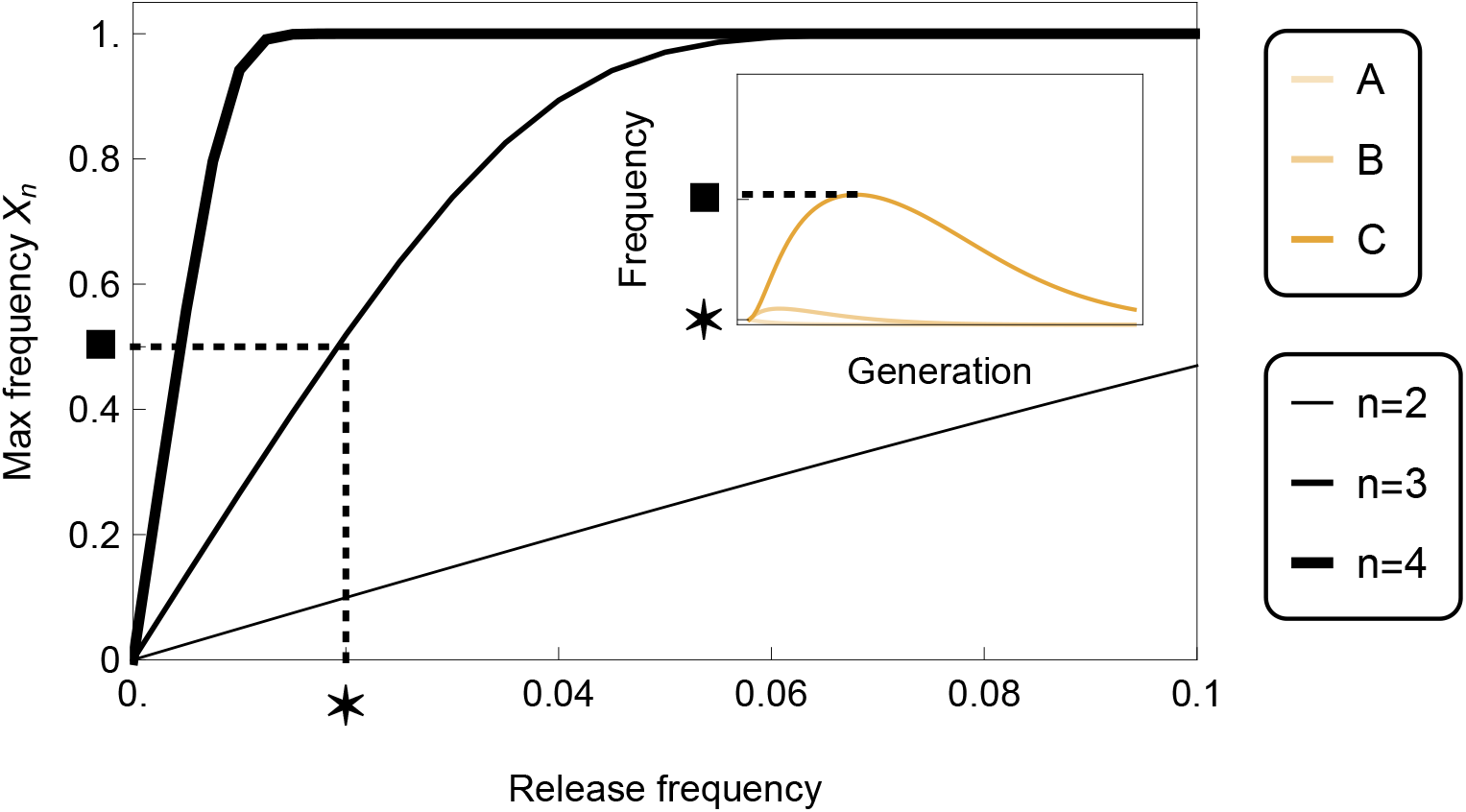
Maximum frequency of the last element in an *n* length daisy-chain construct like locus **A** and **B** in Table 1 in a single population. All elements of the daisy-chain carry a drive load *s*_*d*_ = 0.05. The frequency of the n^th^ transgenic allele is denoted by *X*_*n*_. Other parameters are *δ* = 1, *r* = 0.5.

### 7.2 Mating tables

Here we display the mating tables associated with the 4-locus daisy-quorum system that we are exploring in the main text (Table 1). Table S1 gives the mating table for the dynamics at loci **A** and **B** (the daisy chain component). Table S2 provides the mating table for the dynamics at loci **C** and **D** (the underdominance component).

### 7.3 Alternative payload fitness implementation

In the main text we focus on a dominant payload expression (Table 2), because under this fitness regime, fixation of the *CD* haplotype is locally stable. When we assume different payload regimes (multiplicative dominance or recessive effects on individual fitness), then fixation of *CD* haplotype is no longer a stable equilibrium, as described below.

**Table S1:** Mating table for locus **A** and **B**, illustrating the gametes that come together to make a diploid individual (first two columns), their fitness (third column), at birth frequency (fourth column), and the gametes they produce (fifth-eighth columns)

| Gamete 1 | Gamete 2 | Fitness | Freq | Gametes produced |  |  |  |
| --- | --- | --- | --- | --- | --- | --- | --- |
|  |  |  |  | *ab* | *aB* | *Ab* | *AB* |
| *ab* | *ab* | 1 | $X_{ab}^2$ | 1 | | | |
| *ab* | *aB* | $(1 - s_d)$ | $2X_{ab}X_{aB}$ | $\frac{1}{2}$ | $\frac{1}{2}$ | | |
| *ab* | *Ab* | $(1 - s_d)$ | $2X_{ab}X_{Ab}$ | $\frac{1}{2}$ | | $\frac{1}{2}$ | |
| *ab* | *AB* | $(1 - s_d)^2$ | $2X_{ab}X_{AB}$ | $\frac{1}{2}(1 - \delta)(1 - r)$ | $\frac{1}{2}(\delta + (1 - \delta)r)$ | $\frac{1}{2}(1 - \delta)r$ | $\frac{1}{2}(1 - (1 - \delta)r)$ |
| *aB* | *aB* | $(1 - s_d)^2$ | $X_{aB}^2$ | | 1 | | |
| *aB* | *Ab* | $(1 - s_d)^2$ | $2X_{aB}X_{Ab}$ | $\frac{1}{2}(1 - \delta)r$ | $\frac{1}{2}(1 - (1 - \delta)r)$ | $\frac{1}{2}(1 - \delta)(1 - r)$ | $\frac{1}{2}(\delta + (1 - \delta)r)$ |
| *aB* | *AB* | $(1 - s_d)^3$ | $2X_{aB}X_{AB}$ | | $\frac{1}{2}$ | $\frac{1}{2}$ | |
| *Ab* | *Ab* | $(1 - s_d)^2$ | $X_{Ab}^2$ | | | 1 | |
| *Ab* | *AB* | $(1 - s_d)^3$ | $2X_{Ab}X_{AB}$ | | | $\frac{1}{2}(1 - \delta)$ | $\frac{1}{2}(1 + \delta)$ |
| *AB* | *AB* | $(1 - s_d)^4$ | $X_{AB}^2$ | | | | 1 |

**Table S2:** Mating table for locus **C** and **D**, illustrating the gametes that come together to make a diploid individual (first two columns), their fitness (third column), at birth frequency (fourth column), and the gametes they produce (fifth-eighth columns)

| Gamete 1 | Gamete 2 | Fitness | Freq | Gametes produced |  |  |  |
| --- | --- | --- | --- | --- | --- | --- | --- |
|  |  |  |  | *cd* | *cD* | *Cd* | *CD* |
| *cd* | *cd* | 1 | $X_{cd}^2$ | 1 | | | |
| *cd* | *cD* | $(1 - s_t)(1 - s_p)$ | $2X_{cd}X_{cD}$ | $\frac{1}{2}$ | $\frac{1}{2}$ | | |
| *cd* | *Cd* | $(1 - s_t)(1 - s_p)$ | $2X_{cd}X_{Cd}$ | $\frac{1}{2}$ | | $\frac{1}{2}$ | |
| *cd* | *CD* | $(1 - s_p)$ | $2X_{cd}X_{CD}$ | $\frac{1}{2}(1 - r)$ | $\frac{1}{2}r$ | $\frac{1}{2}r$ | $\frac{1}{2}(1 - r)$ |
| *cD* | *cD* | $(1 - s_t)(1 - s_p)$ | $X_{cD}^2$ | | 1 | | |
| *cD* | *Cd* | $(1 - s_p)$ | $2X_{cD}X_{Cd}$ | $\frac{1}{2}r$ | $\frac{1}{2}(1 - r)$ | $\frac{1}{2}(1 - r)$ | $\frac{1}{2}r$ |
| *cD* | *CD* | $(1 - s_p)$ | $2X_{cD}X_{CD}$ | | $\frac{1}{2}$ | $\frac{1}{2}$ | |
| *Cd* | *Cd* | $(1 - s_t)(1 - s_p)$ | $X_{Cd}^2$ | | | 1 | |
| *Cd* | *CD* | $(1 - s_p)$ | $2X_{Cd}X_{CD}$ | | | $\frac{1}{2}$ | $\frac{1}{2}$ |
| *CD* | *CD* | $(1 - s_p)$ | $X_{CD}^2$ | | | | 1 |

#### Multiplicative dominance of payload

Table S3 shows the individual fitness under a multiplicative dominance regime for the payload.

**Table S3:** Payload (*s*_*p*_) is multiplicative within and between loci

|  | *cd* | *cD* | *Cd* | *CD* |
| --- | --- | --- | --- | --- |
| *cd* | 1 | $(1 - s_t)(1 - s_p)^{1/4}$ | $(1 - s_t)(1 - s_p)^{1/4}$ | $(1 - s_p)^{1/2}$ |
| *cD* | $(1 - s_t)(1 - s_p)^{1/4}$ | $(1 - s_t)(1 - s_p)^{1/2}$ | $(1 - s_p)^{1/2}$ | $(1 - s_p)^{3/4}$ |
| *Cd* | $(1 - s_t)(1 - s_p)^{1/4}$ | $(1 - s_p)^{1/2}$ | $(1 - s_t)(1 - s_p)^{1/2}$ | $(1 - s_p)^{3/4}$ |
| *CD* | $(1 - s_p)^{1/2}$ | $(1 - s_p)^{3/4}$ | $(1 - s_p)^{3/4}$ | $1 - s_p$ |

The governing equations for the underdominance component with multiplicative payload selection in Table S3 are:

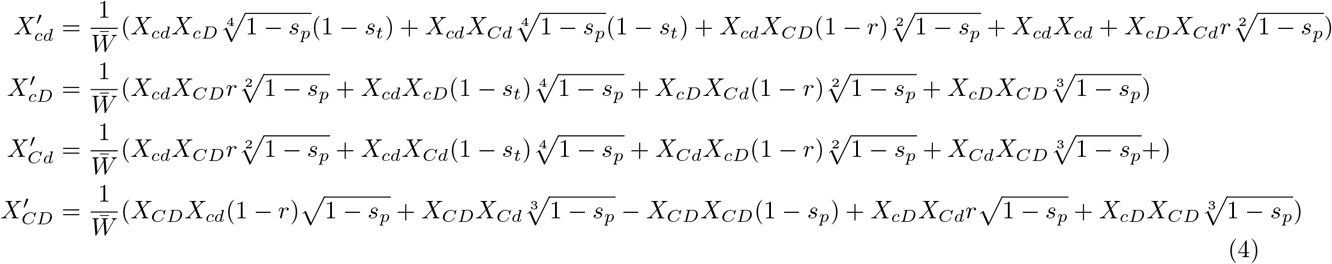

**Figure S2:**
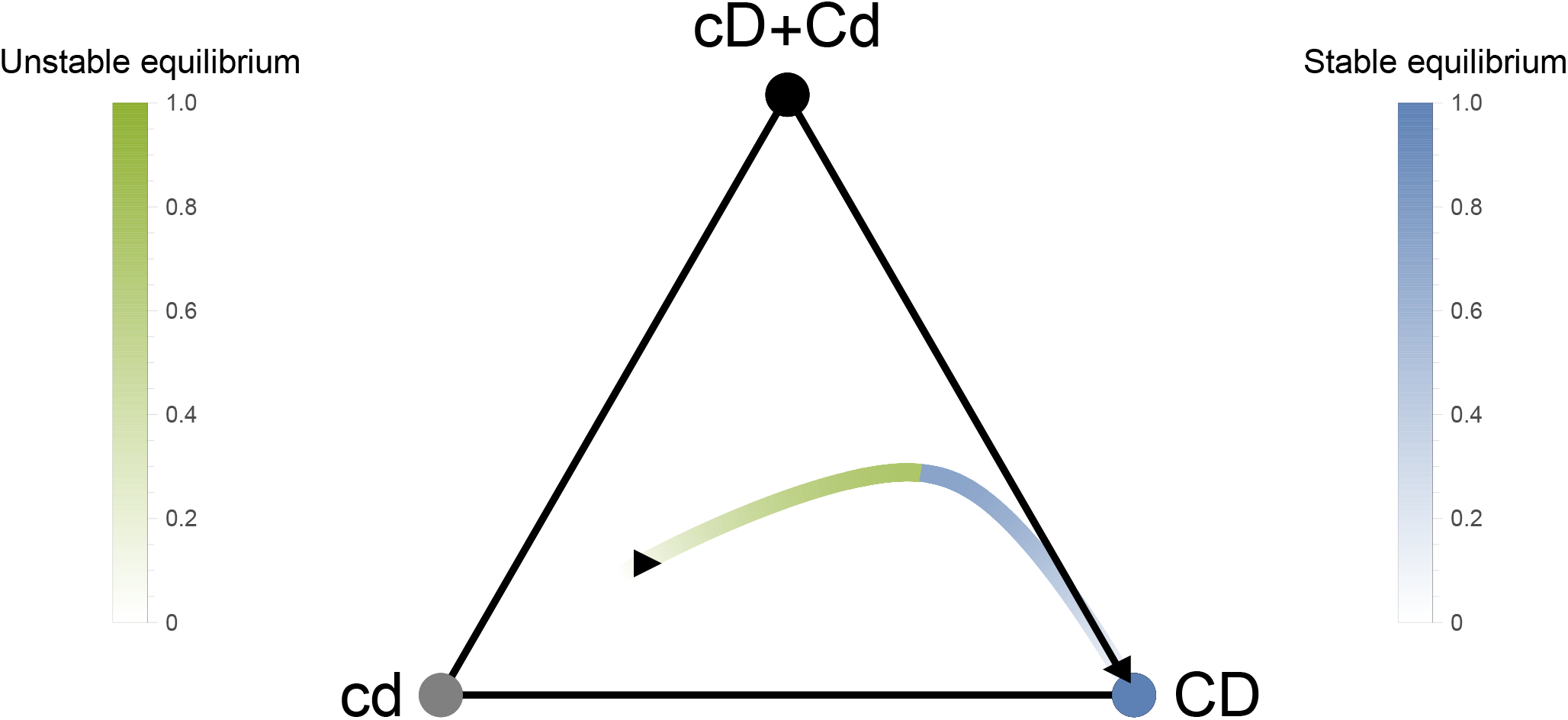
Equilibria of the multiplicative fitness regime of Table S3. The parametric lines indicate the internal unstable (green) and stable (blue) equilibria for different payload values *s*_*p*_. The toxin load *s*_*t*_ = 1 and 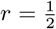. For a given payload, *s*_*p*_, starting to the right of the green point will lead the system to approach the blue point with the same shade (*s*_*p*_ value).

Analysis of the equilibria of this dynamical system reveals two internal equilibria, plus *X*_*cd*_ = 1, which is stable, and *X*_*CD*_ = 1, which is unstable. In Figure S2 we illustrate the location of the various stable and unstable equilibria. As the payload increases, the stable and unstable equilibrium shift towards each other. This means that a drive is harder to invade and will spread to a lower frequency when the payload increases in strength.

The same invasion analysis as in section 3.3 reveals that the conditions for the drive rate *δ*_*c*_ needed for the transgenic alleles *C* and *D* to spread are:

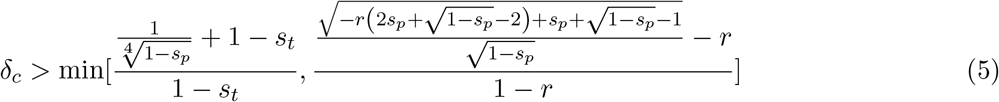

This shows that the maximum payload that can be invaded by a two-locus underdominance system with multiplicative payload is *s*_*p*_ = 15/16 with maximal drive rate *δ* = 1 and *s*_*t*_ ≈ 0.

#### Recessive payload

Table S4 presents the relative fitness of individuals when the payload is recessive, by which we mean an individual needs two copies of *C* and two copies of *D* alleles before expressing the payload.

**Table S4:** Payload (*s*_*p*_) is recessive

| | $cd$ | $cD$ | $Cd$ | $CD$ |
| --- | --- | --- | --- | --- |
| $cd$ | 1 | $1 - s_t$ | $1 - s_t$ | 1 |
| $cD$ | $1 - s_t$ | $1 - s_t$ | 1 | 1 |
| $Cd$ | $1 - s_t$ | 1 | $1 - s_t$ | 1 |
| $CD$ | 1 | 1 | 1 | $1 - s_p$ |

The governing equations for the underdominance component with a recessive payload are:

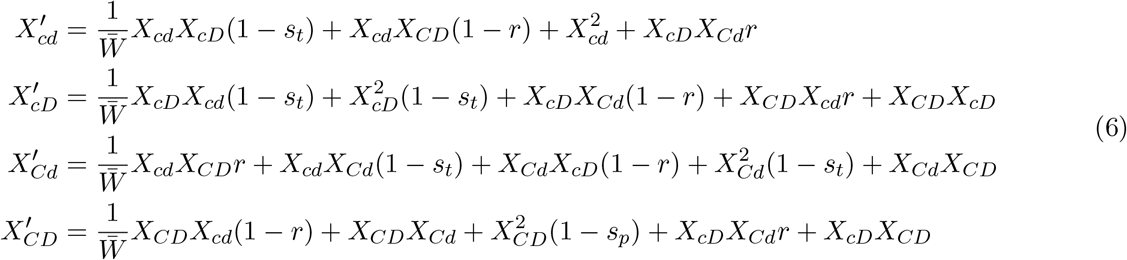

In Figure S3 we illustrate the location of the equilibria of this model for *s*_*t*_ = 1.0 and *s*_*t*_ = 0.1. We see that similar to the analysis in the main text for a dominant payload, a reduction in toxin efficiency brings the equilibrium further from the wildtype (Figure S3 (a,b)).

The results of the invasion analysis show that this construct can only invade when:

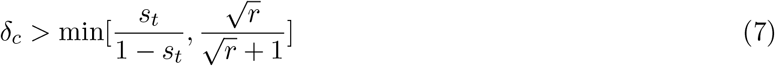

### 7.4 Individual-based simulations

Individual-based simulations (implemented in *C*^++^) were developed to explore whether our results hold for finite population sizes. We use the same fitness regime as in the main text (dominant payload expression). We implement the following life cycle in an isolated population:

1. Calculate the expected frequency of genotypes predicted under the deterministic model.
2. Calculate mean fitness, 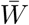, and draw from a Poisson random variate with rate parameter 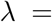 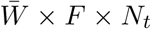 the number of offspring in the next generation 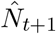. Here, *F* represents the average number of offspring per individual. The total number of offspring in the next generation was then set to 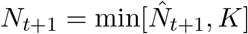.
3. Draw *N*_*t*+1_ offspring genotypes from a multinomial distribution with the deterministic predicted genotype frequencies.

**Figure S3:**
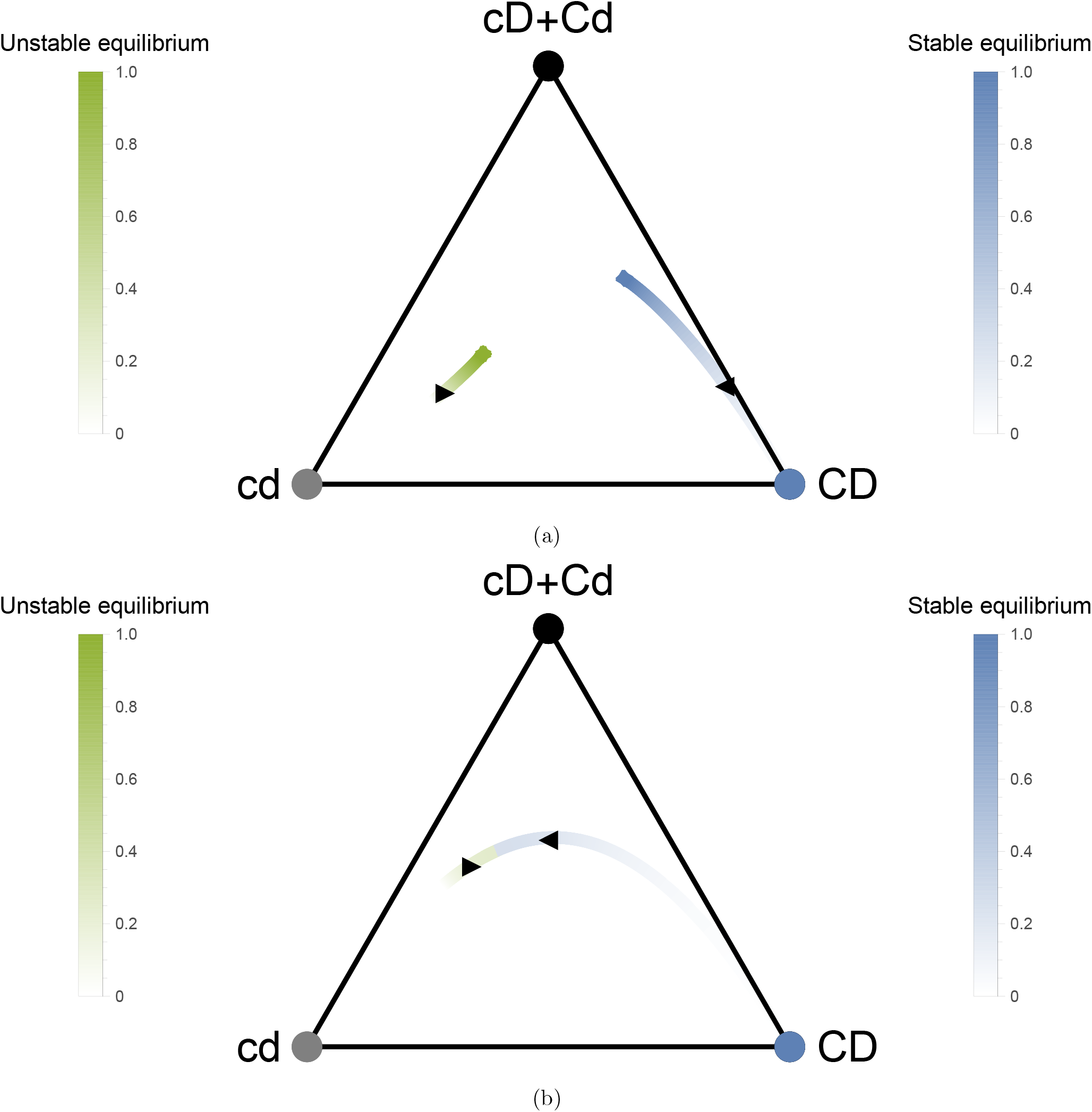
Equilibria of the recessive fitness regime of Table S4 for (a) *s*_*t*_ = 1.0 and (b) *s*_*t*_ = 0.1. The parametric lines indicate the internal unstable (green) and stable (blue) equilibria for different payload values *s*_*p*_. The toxin load *s*_*t*_ = 1 and 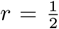. For a given payload, *s*_*p*_, starting to the right of the green point will lead the system to approach the blue point with the same shade (*s*_*p*_ value).

**Figure S4:**
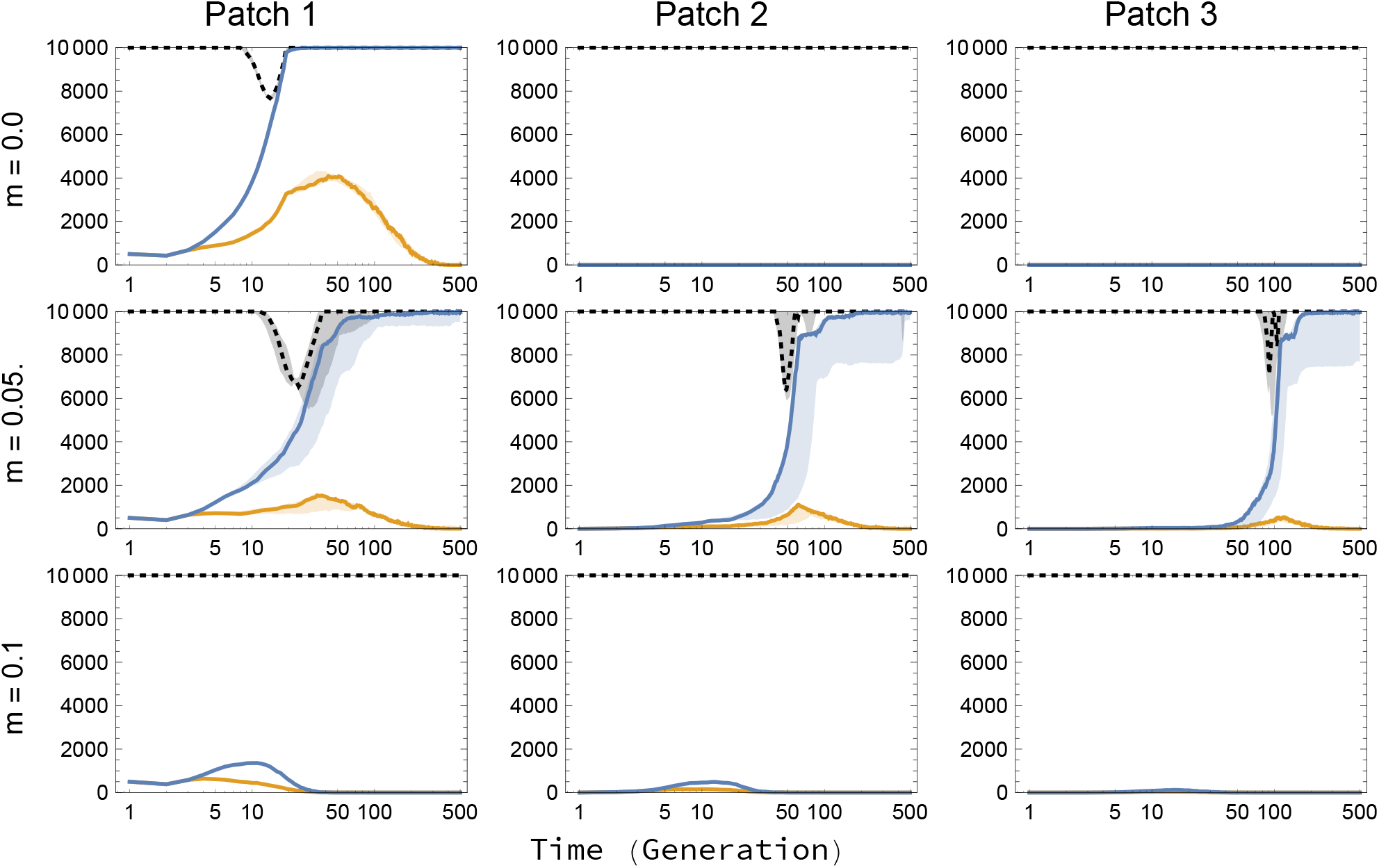
Finite population demographic dynamics. Black dotted line is the total population size at time *t*, blue solid line represents the number of individuals carrying the payload allele and orange solid line represents the number of individuals carrying the *B* allele. In this case, fertility is set to *F* = 1.2 and the payload to *s*_*p*_ = 0.1, so that patches can persist even if the underdominance construct fixes. Each patch has a carrying capacity of *K* = 10000. Rows have a migration probability of 0, 0.05, 0.1 from top to bottom. Drive is released on the left. Results are shown for 50 replicates. The solid line represents the median of the replicates, and the shaded region the first and third quantiles.

We repeat these steps for *t*_*max*_ generations and show results in Figure S4 and S5 for *F* = 1.2 and *F* = 1.05, respectively. In Figure S4, with *F* = 1.2 and a payload of *s*_*p*_ = 0.1, the average number of offspring per individual carrying the payload remains above unity: *λ* = 0.9 × 1.2 = 1.08. However, the toxin load *s*_*t*_ = 0.9 causes a dip in population size (black dashed line) as the payload increases in frequency. Once the payload has fixed, the toxin load will not be expressed, and the population recovers to its carrying capacity. Figure S5 illustrates dynamics for *F* = 1.05. Here *λ* = 0.9 × 1.05 = 0.945, which is below unity, meaning that individuals carrying the payload will not be able to replace themselves on average. We see that in this scenario the payload for *m* = 0.05 does not go to fixation, unlike Figure S4. This is because the population size is more severely affected by the payload. As the population declines, proportionally more migrants enter the patch than when the population is large. This proportionally high inflow of wildtype individuals prevents fixation of the payload and eventually helps the population recover to its carrying capacity with only wildtype individuals.

**Figure S5:**
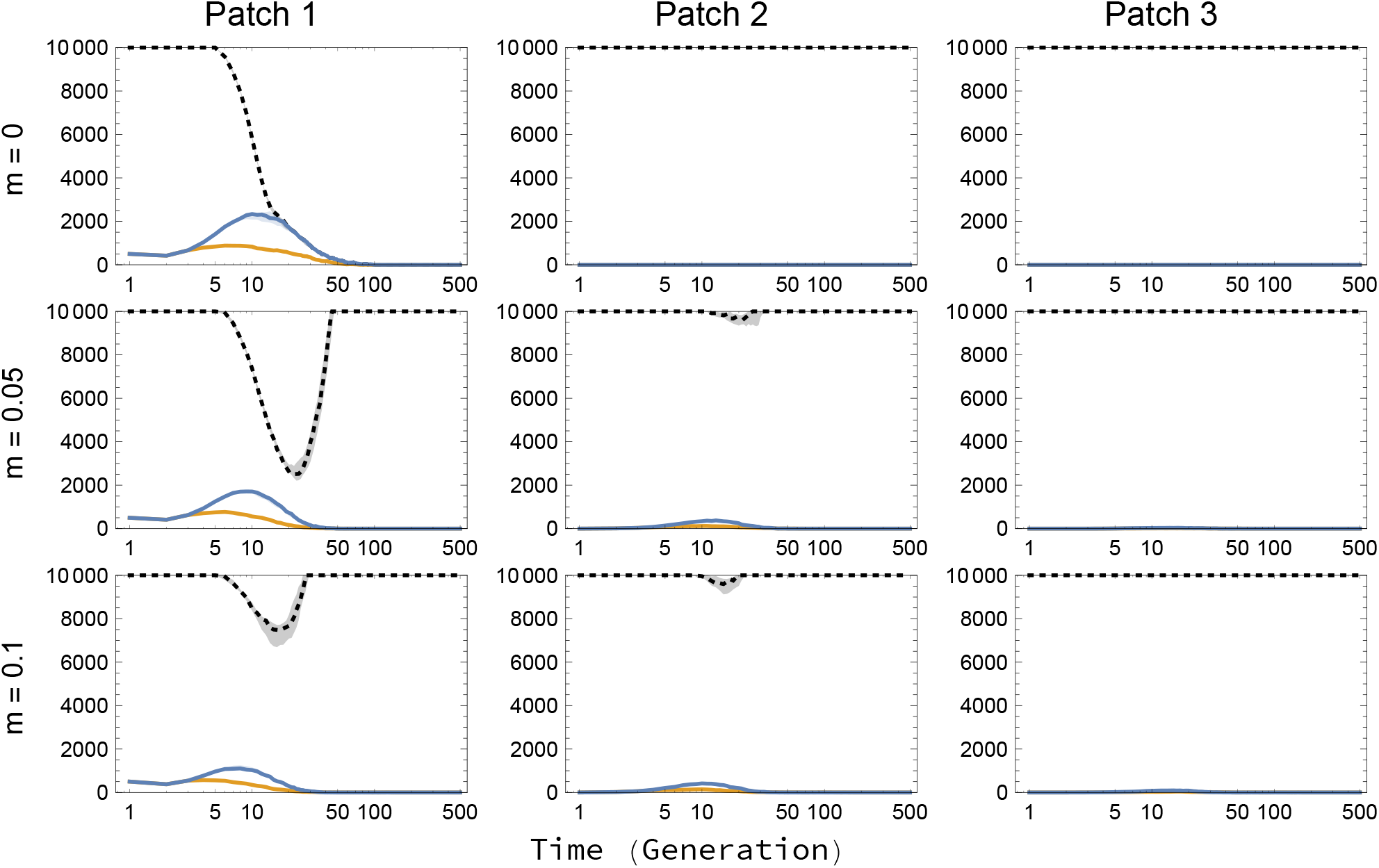
Finite population demographic dynamics. Black dotted line is the total population size at time *t*, blue solid line represents the number of individuals carrying the payload allele and orange solid line represents the number of individuals carrying the *B* allele. In this case, fertility is set to *F* = 1.05 and the payload to *s*_*p*_ = 0.1, so that patches are driven extinct if the payload fixes after which recolonization by the wildtype eventually occurs. Each patch has a carrying capacity of *K* = 10000. Rows have a migration probability of 0, 0.05, 0.1 from top to bottom. Drive is released on the left. Results are shown for 50 replicates. The solid line represents the median of the replicates, and the shaded region the first and third quantiles.

In our deterministic analysis of spread in Figures 4 and 5, we noted that a decrease in the toxin load *s*_*t*_ can help prevent spillover of the construct by shifting the separatrix further from the wildtype state (Figure 2). However, decreasing the toxic load in order to shift the separatrix comes at a cost in terms of persistence of the construct in finite population sizes. In the limit as the toxic load goes to 0, the valley in Figure 1 (panel B) disappears, which results in directional selection towards the wildtype. Lowering the toxin load to center the separatrix (the location of the valley) will simultaneously reduce the depth of the fitness valley. Such a shallow fitness landscape means that the effects of drift in finite population sizes can influence the dynamics significantly.

**Figure S6:**
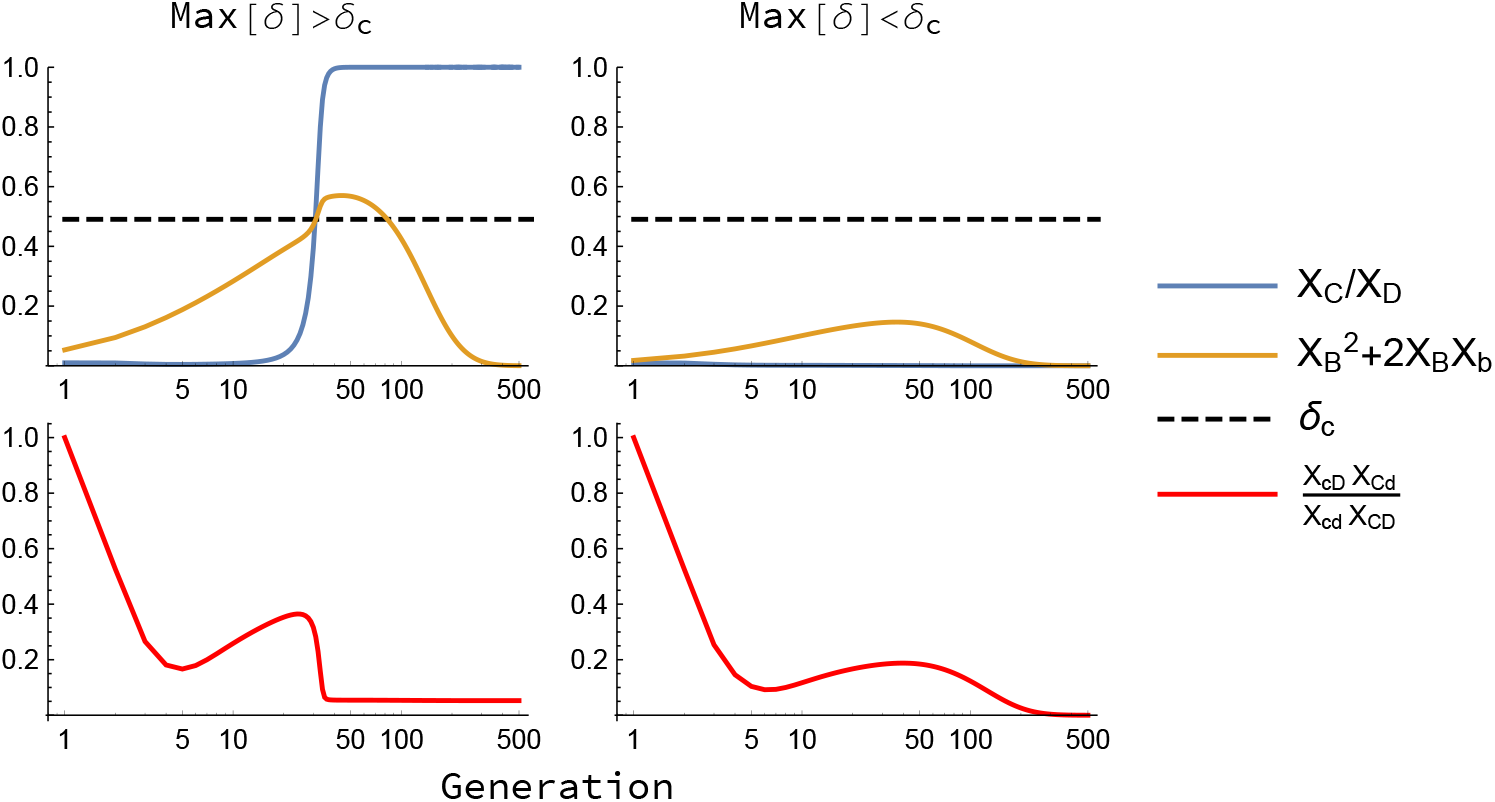
*δ* = 0.9, *r* = 0.5, *s*_*p*_ = 0.1, *s*_*t*_ = 0.9, *f*_*drive*_ = 0.01|0.03 and *f*_*EUD*_ = 0.01

### 7.5 Invasion analysis

Our invasion analysis relies on there not being any linkage disequilibrium between the loci involved in the daisy chain and the underdominance component. In Figure S6 we illustrate the critical drive threshold above which the *X*_*cd*_ = 1 equilibrium becomes unstable assuming there is no linkage disequilibrium between the loci. The left column shows a starting frequency of the *CD* alleles at *f*_*CD*_ = 0.01 and the right column for *f*_*CD*_ = 0.03. We initiate the simulations without linkage disequilibrium between the **AB** and **CD** loci. The initial frequency for *CD* is *f*_*CD*_ = 0.01. The top graphs show the dynamics of the simulation, where the left column illustrates a scenario where the drive strength *δ* exceeds the critical threshold *δ*_*c*_ while the right does not. The bottom graphs show that significant linkage disequilibrium builds up soon after the simulations start. As a consequence of these genetic associations, the threshold drive does not precisely predict when spread will occur. For example, in the top middle panel, the drive frequency (orange) rises above the threshold (dashed line) but the underdominance construct fails to spread to fixation. Nevertheless, equation (3) provides an approximate guide to when the underdominance can (left) and will not (right) spread.

## Notes

### Competing Interest Statement

The authors have declared no competing interest.

